# An Msp1-Protease Chimera Captures Transient AAA+ Interactions and Unveils Ost4 Mislocalization Errors

**DOI:** 10.1101/2025.03.31.646376

**Authors:** Deepika Gaur, Brian Acquaviva, Matthew L. Wohlever

## Abstract

Membrane protein homeostasis (proteostasis) is essential for maintaining the integrity of eukaryotic organelles. Msp1 is a membrane anchored AAA+ (ATPase Associated with cellular Activities) protein that maintains mitochondrial proteostasis by extracting aberrant proteins from the outer mitochondrial membrane. A comprehensive understanding of the physiological roles of Msp1 has been hindered because AAA+ proteins interact with substrates transiently and common strategies to stabilize this interaction lead to undesirable mitochondrial phenotypes. To circumvent these drawbacks, we fused catalytically active Msp1 to the inactivated protease domain of the AAA+ protease Yme1. The resulting chimera sequesters substrates in the catalytically inactive degradation chamber formed by the protease domain. We performed mass spectrometry analysis with the Msp1-protease chimera and identified the signal anchored protein Ost4 as a novel Msp1 substrate. Topology experiments show that Ost4 adopts mixed orientations when mislocalized to mitochondria and that Msp1 extracts mislocalized Ost4 regardless of orientation. Together, this work develops new tools for capturing transient interactions with AAA+ proteins, identifies new Msp1 substrates, and shows a surprising error in targeting of Ost4.

## Introduction

Mitochondria contain >1000 proteins, over 99% of which are nuclear encoded and need to be transported into or across the outer mitochondrial membrane (OMM)(Brave *et al*, 2023; Calvo *et al*, 2016; Busch *et al*, 2023; Muthukumar *et al*, 2024). Protein targeting is imperfect; therefore cells are heavily reliant on quality control mechanisms to maintain mitochondrial proteostasis(Krämer *et al*, 2023; Itakura *et al*, 2016; Hansen *et al*, 2018; Jiang, 2020; Song *et al*, 2021; Song & Becker, 2022). Failure to maintain mitochondrial proteostasis is implicated in various diseases such as neurodegeneration, cancer, and aging(Benaroya, 2024; Uoselis *et al*, 2023). An essential aspect of mitochondrial proteostasis is the selective removal of proteins from the mitochondrial membrane(Wilson *et al*, 2024; Uoselis *et al*, 2023; Mukhtar *et al*, 2023; Yamano *et al*, 2023; Rödl & Herrmann, 2023). This critical process is driven by AAA+ (ATPases Associated with cellular Activities) molecular motors, which utilize ATP to perform the thermodynamically unfavorable task of extracting hydrophobic substrates from a lipid bilayer(Glynn *et al*, 2020; Wohlever *et al*, 2017; Smith *et al*, 2024; Lin *et al*, 2022; Hanson & Whiteheart, 2005; Fresenius *et al*, 2023).

One such AAA+ protein is Msp1, which localizes to the OMM and peroxisomes and promotes mitochondrial proteostasis by removing mislocalized tail-anchored (TA) proteins (Wang & Walter, 2020; Fresenius & Wohlever, 2019; Matsumoto & Endo, 2022; Weir *et al*, 2017; Castanzo *et al*, 2020; Matsumoto *et al*, 2022). In addition to a role in extracting TA proteins, Msp1 is recruited to the TOM complex to remove proteins that stall during translocation across the OMM(Weidberg & Amon, 2018a; Kim *et al*, 2024). Loss of Msp1, especially when combined with a compromised GET pathway, leads to severe mitochondrial defects including failures in oxidative phosphorylation, loss of mitochondrial DNA, and mitochondrial fragmentation (Chen *et al*, 2014; Okreglak & Walter, 2014). In addition to the aforementioned quality control roles, the metazoan homolog ATAD1 has been shown to regulate apoptosis and AMPA receptor internalization(Winter *et al*, 2022; Zhang *et al*, 2011).

Despite clear physiological significance, the role of Msp1 in mitochondrial membrane proteostasis is incompletely defined. To date, only a handful of bona fide Msp1 substrates have been identified(Chen *et al*, 2014; He *et al*, 2023; Li *et al*, 2019; Wang *et al*, 2022). As almost all Msp1 substrates are tail-anchored proteins with a short C-terminal domain in the intermembrane space (IMS), it is currently unclear if Msp1 can extract membrane proteins with the opposite topology of the N-terminus facing the IMS.

A major obstacle to identifying Msp1 substrates is that the interaction of AAA+ proteins with substrates is transient. A common strategy is to use the E to Q Walker B mutation to inhibit ATP hydrolysis and thus generate a substrate trap (Wang *et al*, 2020; Chen *et al*, 2014). An obvious downside to this approach is that the Walker B mutation inhibits AAA+ protein activity, which can lead to altered cell physiology, and therefore altered substrate profiles.

To address this challenge, we adapted a technique that has been used to identify substrates for the AAA+ proteases FtsH and ClpXP which involves creating a “protease trap” mutant with an active ATPase domain (Flynn *et al*, 2003; Westphal *et al*, 2012). AAA+ proteases sequester the protease active sites in a degradation chamber that can only be accessed by translocation through the axial pore of the AAA+ unfoldase (Sauer *et al*, 2022) (**Figure 1A**). Mutation of the catalytically active residues in the protease domain results in substrates becoming trapped in the degradation chamber because only the degradation products (small peptides) can freely diffuse out of the degradation chamber.

**Figure 1:**
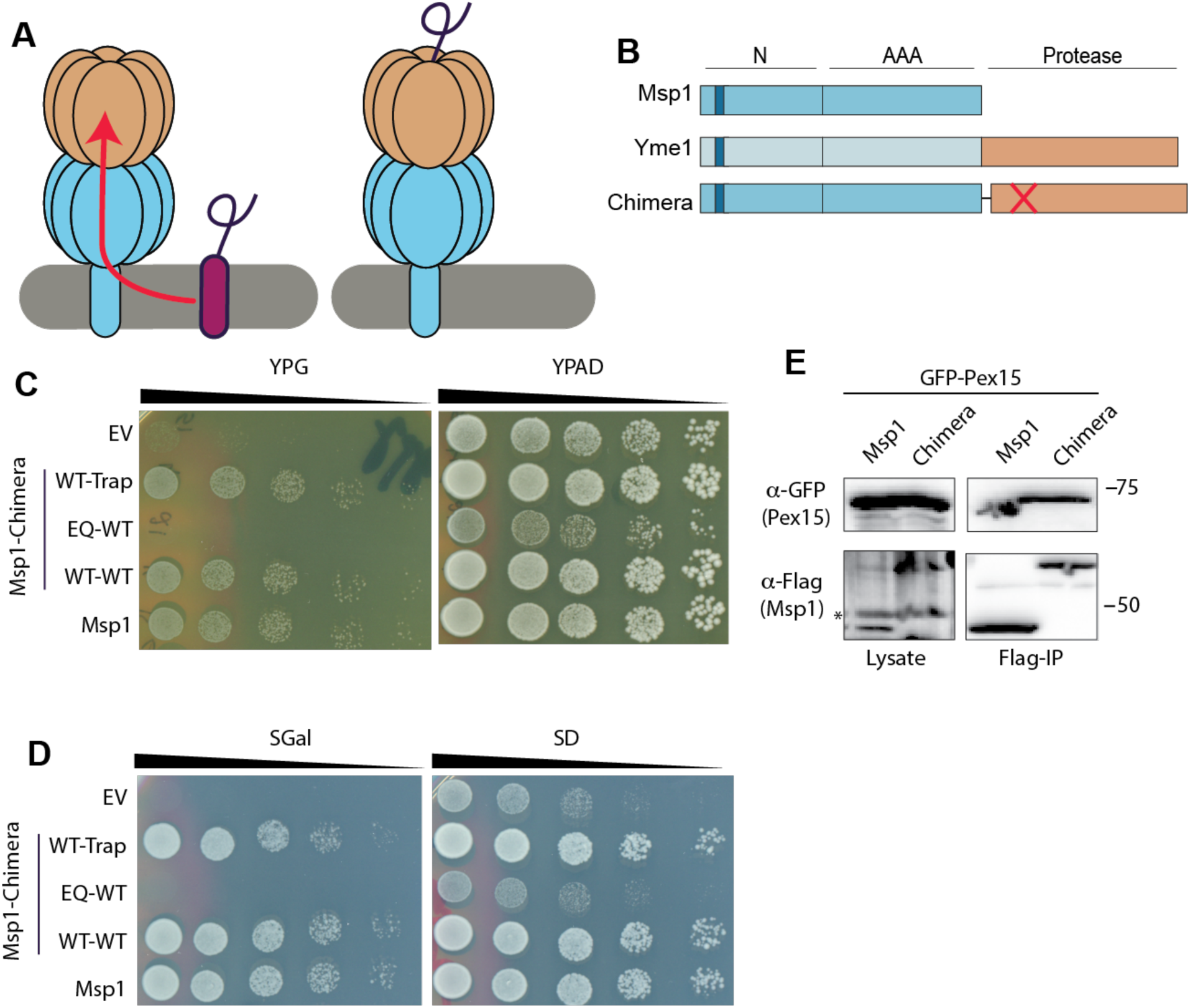
Development of the Msp1-protease chimera. A) Diagram of the protease chimera strategy B) Domain diagram of the protease chimera. The protease inactivating H540Y mutation is indicated by the red X. C) The protease chimera can rescue growth of *Msp1Δ, Get3Δ* yeast on glycerol. Rescue depends on ATPase activity, but not protease activity. WT-Trap is a chimera with WT Msp1 fused to the Yme1 protease domain with the H540Y mutation. EQ-WT is a chimera with the catalytically inactive E193Q mutation in Msp1 fused to the wild type Yme1 protease domain. D) The protease chimera can alleviate toxicity of Pex15 overexpression in *Msp1Δ*, *Get3Δ* yeast. E) The protease chimera interacts with the known Msp1 substrate Pex15 better than WT Msp1. *Msp1Δ*, *Get3Δ* cells. Non-specific band in lysate marked by *.

We have adapted this technique to work with Msp1 by fusing the AAA+ domain to the catalytically inactive protease domain from the membrane anchored protease Yme1. The resulting chimera retains full Msp1 functionality and can trap substrates more efficiently that WT Msp1. Using our chimera, we identified and validated that the ER signal anchored protein Ost4 as a new Msp1 substrate. Surprisingly, we observed that mislocalized Ost4 adopts mixed topologies, with either the N or C-terminus facing the IMS. We demonstrate that Msp1 robustly extracts Ost4 regardless of orientation. Together, our results identify new Msp1 substrates, demonstrate that Ost4 topology is lost upon mistargeting to the OMM, establish that Msp1 can extract substrates with both tail-anchored and signal anchored orientations, and develop a generalizable technique for identifying transient interactors with AAA+ proteins.

## Results

### Development of catalytically active Msp1 substrate trap

We generated a protease-chimera by using an eight-residue serine glycine linker to fuse the protease domain of *S. cerevisiae* Yme1 onto the C-terminus of Msp1 (**Figure 1B**). To function as a substrate trap, the protease chimera should retain the functionality of WT Msp1 while also showing an enhanced ability to trap substrates. We therefore defined the following criteria for a fully functional protease chimera: 1) rescue the growth defect of *Msp1Δ, Get3Δ* yeast on glycerol in an ATPase dependent, but protease independent manner, 2) resolve the toxicity from overexpression of the Msp1 substrate Pex15, and 3) interact with the known Msp1 substrate Pex15 better than WT Msp1.

To assess if the protease chimera retains Msp1 functionality, we first assayed the ability to rescue the growth defect of *Msp1Δ, Get3Δ* yeast on glycerol. *Get3* is a key component of the Guided Entry of Tail-anchored proteins (GET pathway), which targets tail-anchored (TA) proteins to the endoplasmic reticulum. Loss of Get3 results in the mislocalization of ER TA proteins to the mitochondria, which requires Msp1 to clear the mislocalized proteins. The accumulation of mislocalized proteins on the mitochondria in *Msp1Δ, Get3Δ* yeast results in a loss of oxidative phosphorylation, which can be assayed by growth on a non-fermentable carbon source, such as glycerol(Chen *et al*, 2014). The Msp1-protease chimera rescued growth better than the empty plasmid negative control. Assays with the ATPase deficient E193Q mutant and protease deficient H540Y mutant demonstrate that complementation depends on ATPase activity, but not protease activity (**Figure 1C**).

We next tested the ability of the protease chimera to resolve the toxicity from overexpression of the Msp1 substrate Pex15(Chen *et al*, 2014). *Msp1Δ, Get3Δ* yeast were complemented with the protease chimera and Pex15 was overexpressed on a plasmid with a Gal promoter. The protease chimera rescued the phenotype. Once again, rescue depended on ATPase activity, but not protease activity (**Figure 1D**).

We then tested the ability of the protease chimera to interact with the known Msp1 substrate Pex15(Chen *et al*, 2014). We appended a 1x Flag tag to the C-terminus of WT Msp1 or the protease chimera with an inactivated protease domain and then performed an anti-Flag immunoprecipitation (IP). As expected, significantly more Pex15 was present in the elution fraction of with the protease chimera than WT Msp1 (**Figure 1E**). We conclude that the protease chimera retains the functionality of WT Msp1 while also functioning as a substrate trap.

### Mass spectrometry analysis of interactors with Msp1-protease chimera

To identify Msp1 substrates, we generated a *Msp1Δ* yeast strain complemented with the centromeric plasmid pRS315 containing either the Msp1-protease chimera or isolated protease domain as a negative control. Cells were grown to mid-log phase in SD – Leu media, lysed with a bead beater, and solubilized with digitonin before performing an anti-Flag immunoprecipitation (IP). The elution from both samples was trypsin digested and analyzed by LC-MS/MS via a nanoUPLC linked to an Orbitrap Fusion Lumos mass spectrometer. We refer to this as our “standard conditions dataset”.

Previous work demonstrated that Msp1 is recruited to the TOM complex to remove stalled proteins which accumulate during mitochondrial protein import stress (Kim *et al*, 2024; Weidberg & Amon, 2018b). To understand how recruitment to the TOM complex may change the Msp1 substrate profile, we treated our two cell lines with the 20 μM of the ionophore CCCP for 2 h prior to harvesting. CCCP leads to stalling of mitochondrial protein import via the dissipation of the proton gradient. We refer to this as our “CCCP dataset”.

The standard condition dataset included a total of 220 proteins, of which 93 were unique to the protease chimera and 127 were also observed in the protease domain only negative control **(Figure S1A)**. The CCCP dataset contained a total of 196 proteins, of which 56 were unique to the chimera **(Figure S1B)**. There was a combined total of 125 chimera specific hits across both datasets, with 69 observed only in standard conditions, 32 observed only in CCCP conditions, and 24 observed in both conditions **(Figure S1C)**.

To remove non-specific binders from our dataset, we plotted the average spectral counts for each candidate protein that are present in the “CRAPome” of common non-specific binders IP-mass spec datasets(Mellacheruvu *et al*, 2013). We chose to focus our analysis on candidate proteins with a score < 2, which resulted in a combined dataset of 55 proteins **(Figure S1D & E)**.

Gene Ontology (GO) analysis of the filtered dataset showed no significant enrichment for any biological process(Ashburner *et al*, 2000; Aleksander *et al*, 2023). Contrary to our hypothesis, GO analysis of the hits unique to the CCCP dataset showed no enrichment for mitochondrial specific pathways and there were few hits that localize to the inner mitochondrial membrane (IMM) **(Tables S1-S5)**. The lack of enrichment for a specific biological process is consistent with Msp1 acting as a general proteostasis factor, which could allow it to interact with substrates involved in numerous different biological processes.

We next compared the combined hits from both the standard condition and CCCP datasets to previously published work using the E193Q mutant(Chen *et al*, 2014). Surprisingly, after filtering out results based on CRAPome spectral counts, the only proteins in common between our all datasets was Phb1 and the trivial result of Msp1. Previously validated substrates Pex15 and Gos1 were not present in any of the datasets, including the protease only negative controls. Indeed, the combined dataset contained 11 membrane proteins and no tail-anchored proteins. We conclude that while our dataset provides a list of candidate Msp1 substrates, this list is not comprehensive, potentially due to challenges in identifying membrane proteins by mass spectrometry. Future work with the protease chimera will benefit from quantitative mass spectrometry approaches to better define substrate enrichment by the protease chimera as well as alternative mass spec protocols to enable more robust detection of membrane proteins.

### Msp1 interacts with Ost4 and Phb1

As GO analysis provided no guidance for candidate substrates, we decided to manually select a handful of hits for further investigation. To make our search as broad as possible, we selected hits that involved a range of cellular locations and processes, were observed in different datasets, and contained a mixture of soluble and membrane bound proteins (**Figure 2A**).

**Figure 2:**
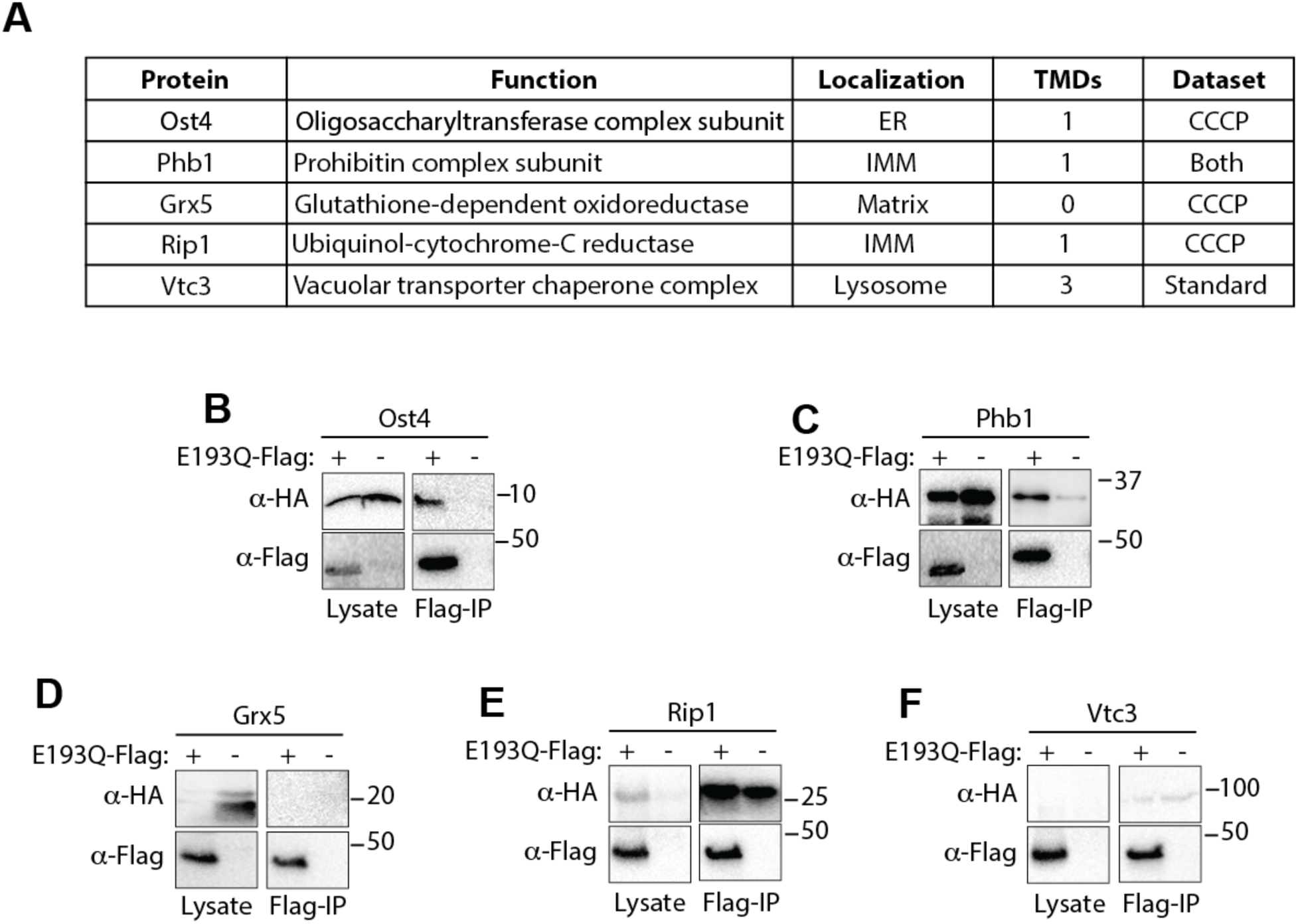
E193Q Msp1 interacts with Ost4 and Phb1. A) Table showing mass spec hits selected for further examination. B-F) *Msp1Δ* cells were transformed with a centromeric plasmid expressing 2x-HA-tagged substrate and a centromeric plasmid expressing E193Q Msp1 with or without a 3x-flag tag. Whole cell lysates were subjected to an anti-flag immunoprecipitation. IP results show that E193Q Msp1 interacts with Ost4 and Phb1.

As an initial test of Msp1 interaction, we cloned a 2x-HA tag onto the substrate and performed an immunoprecipitation in *Msp1Δ* yeast complemented with either 3x-Flag-tagged E193Q Msp1 or untagged E193Q Msp1. We observed robust and specific interaction with Phb1 and Ost4 (**Figure 2B & C)**. The proteins Grx5, Rip1, and Vtc3 either bound non-specifically to the resin or were not detectable by western blot (**Figure 2D-F**). We therefore chose to focus our analysis on Ost4 and Phb1.

Both putative substrates contain a single transmembrane domain, consistent with previously annotated Msp1 substrates. Phb1 is a subunit of the prohibitin complex and has a single N-terminal TMD which anchors it to the inner mitochondrial membrane (IMM)(Steglich *et al*, 1999). Ost4 is subunit of the oligosaccharyl transferase complex(Chi *et al*, 1996; Dumax-Vorzet *et al*, 2013). It is a small protein of only 36 residues and contains a single TMD. Phb1 was observed in both datasets as well as previous studies with E193Q Msp1(Chen *et al*, 2014). Conversely, interaction with Ost4 was only observed in the CCCP dataset and there are no reported genetic or physical interactions between Ost4 and Msp1 on SGD(Cherry *et al*, 1997; Wong *et al*, 2023).

### Msp1 extracts Ost4 from the OMM

To test if Msp1 removes Phb1 and Ost4 from the OMM, we developed luciferase based Msp1 activity assay (**Figure 3A**). This split luciferase assay is a variation of the MitoLuc targeting assay(Needs *et al*, 2023) and involves cloning the 11-residue HiBiT tag onto a substrate and targeting LgBiT to the intermembrane space (IMS). Protein insertion into the OMM is then monitored by luminescence.

**Figure 3:**
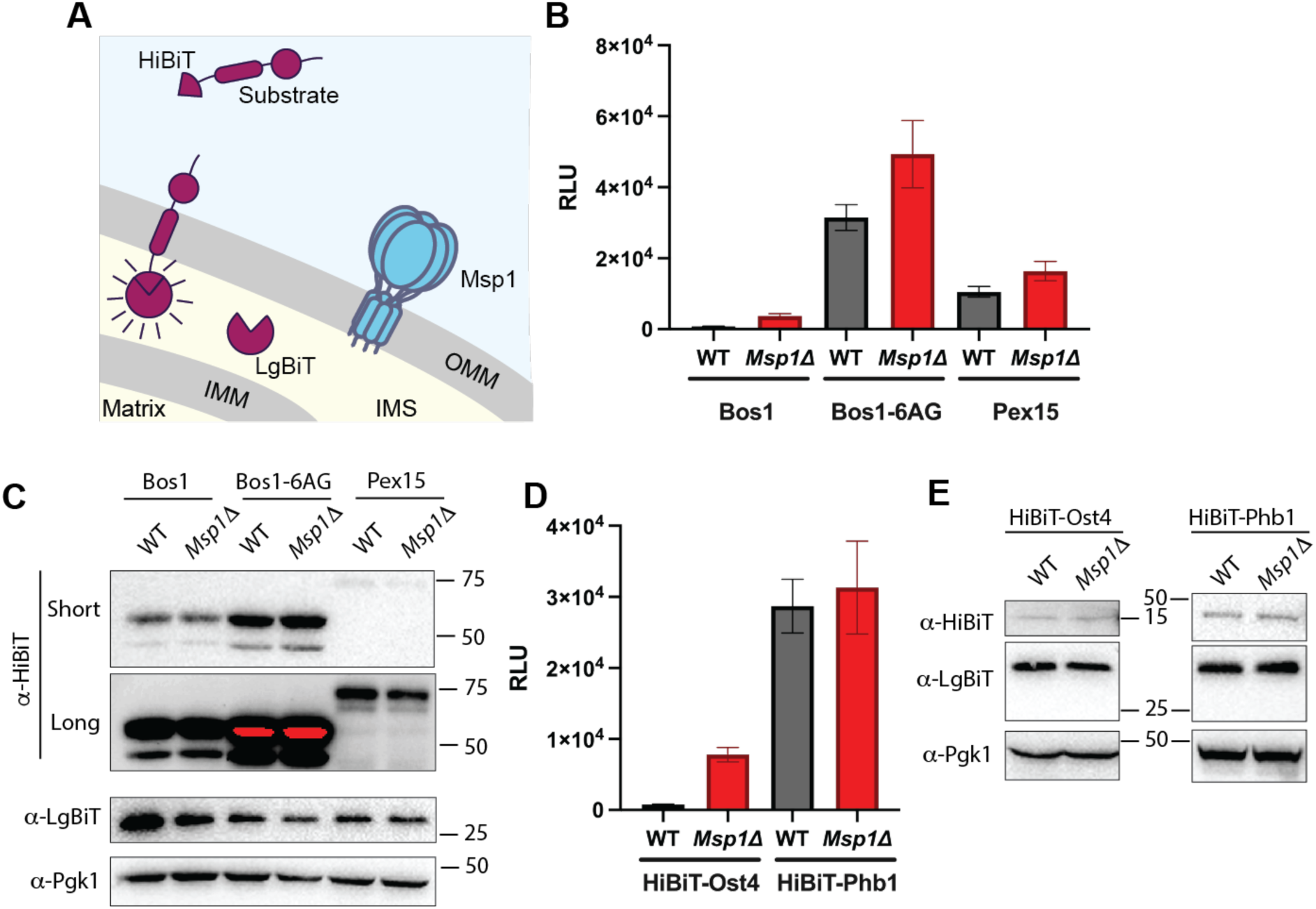
Msp1 removes Ost4 from the OMM. A) Diagram of split luciferase assay to monitor protein insertion and extraction from the OMM. Higher levels of luciferase indicate increased substrate insertion into the OMM. B) Msp1 decreases the OMM concentration of the tail-anchored proteins Bos1-HiBiT, Bos1-6AG-HiBiT, and Pex15-HiBiT. Higher luminescence indicates more HiBiT tagged substrate is inserted in the OMM. Error bars are standard error of the mean from 3 replicates. C) Western blot of the cells from B shows equal expression of HiBiT tagged substrates and IMS localized LgBiT in the WT and *Msp1Δ* cells. The anti-HiBiT blot is shown with both a short and long exposure to allow for visualization of Pex15-HiBiT. D) Msp1 decreases the OMM concentration of HiBiT-Ost4 but has no effect on HiBiT-Phb1. Error bars are standard error of the mean from 3 replicates. E) Western blot of the cells from D shows equal expression of HiBiT tagged substrates and IMS localized LgBiT in the WT and *Msp1Δ* cells.

As proof of principle, we transformed WT and *Msp1Δ* cells with the known Msp1 substrate Pex15-HiBiT, the ER tail-anchored protein Bos1-HiBiT, and the Bos1-6AG-HiBiT mutant, which shows increased mitochondrial targeting (Rao *et al*, 2016). With all three substrates we observed higher levels of luminescence in *Msp1Δ* cells than WT cells suggesting that Msp1 reduces the mitochondrial concentration of tail-anchored proteins (**Figure 3B**). Western blots show equal expression of the substrate and LgBiT in both WT and *Msp1Δ* cells (**Figure 3C**), indicating that the increased signal in *Msp1Δ* cells is due to changes in subcellular localization rather than a difference in LgBiT or HiBiT expression.

With our assay well established, we next tested if Msp1 changes the mitochondrial concentration of the signal anchored proteins HiBiT-Ost4 and HiBiT-Phb1. Similar to known Msp1 substrates, the HiBiT-Ost4 substrate had increased luminesence in *Msp1Δ* cells compared to WT cells (**Figure 3D**). Western blot controls again show equal expression of the HiBiT tagged substrate and LgBiT (**Figure 3E**). We observed no Msp1 dependent difference with HiBiT-Phb1. The split luciferase assay cannot distinguish between Phb1 that is inserted into the OMM and Phb1 that is completely translocated across the OMM en route to the IMM. We suspect that this is swamping out any Msp1 dependent change in signal. We therefore chose to focus only on Ost4 for further analysis.

### Ost4 adopts mixed topology when mislocalized to the OMM and both topologies are extracted by Msp1

Ost4 is a signal anchored membrane protein that is part of the Oligosaccharyl Transferase Complex (OST). Ost4 is annotated as a signal anchored protein, with Cryo-EM structures showing the N-terminus facing the ER lumen(Ramírez *et al*, 2022; Kim *et al*, 2003). At only 36 residues long, the TMD is close enough to the C-terminus that it is likely still in the ribosome exit tunnel when translation finishes, thereby meeting the definition of a tail-anchored protein (Borgese, 2003). We hypothesized that upon mistargeting to the mitochondria, Ost4 could be inserted with either the N- or C-terminus in the intermembrane space (IMS).

To test this hypothesis, we generated the Ost4-HiBiT construct, which will luminesce if it assumes the topology of a tail-anchored protein, with the C-terminus in the IMS. We observed robust luminescence with the Ost4-HiBiT that increases upon deletion of Msp1 (**Figure 4A**). Surprisingly, the Ost4-HiBiT construct showed higher luminescence compared to the HiBiT-Ost4 construct. Western blots show that the increased signal with C-terminally tagged Ost4 likely arises from higher expression levels (**Figure 4B**). It is unclear why there is such a large difference in the stability of N and C-terminally tagged Ost4. Regardless of the difference in expression levels, our data shows that both the N and C-terminus of Ost4 can access the IMS and that the signal intensity increases dramatically when Msp1 is deleted.

**Figure 4:**
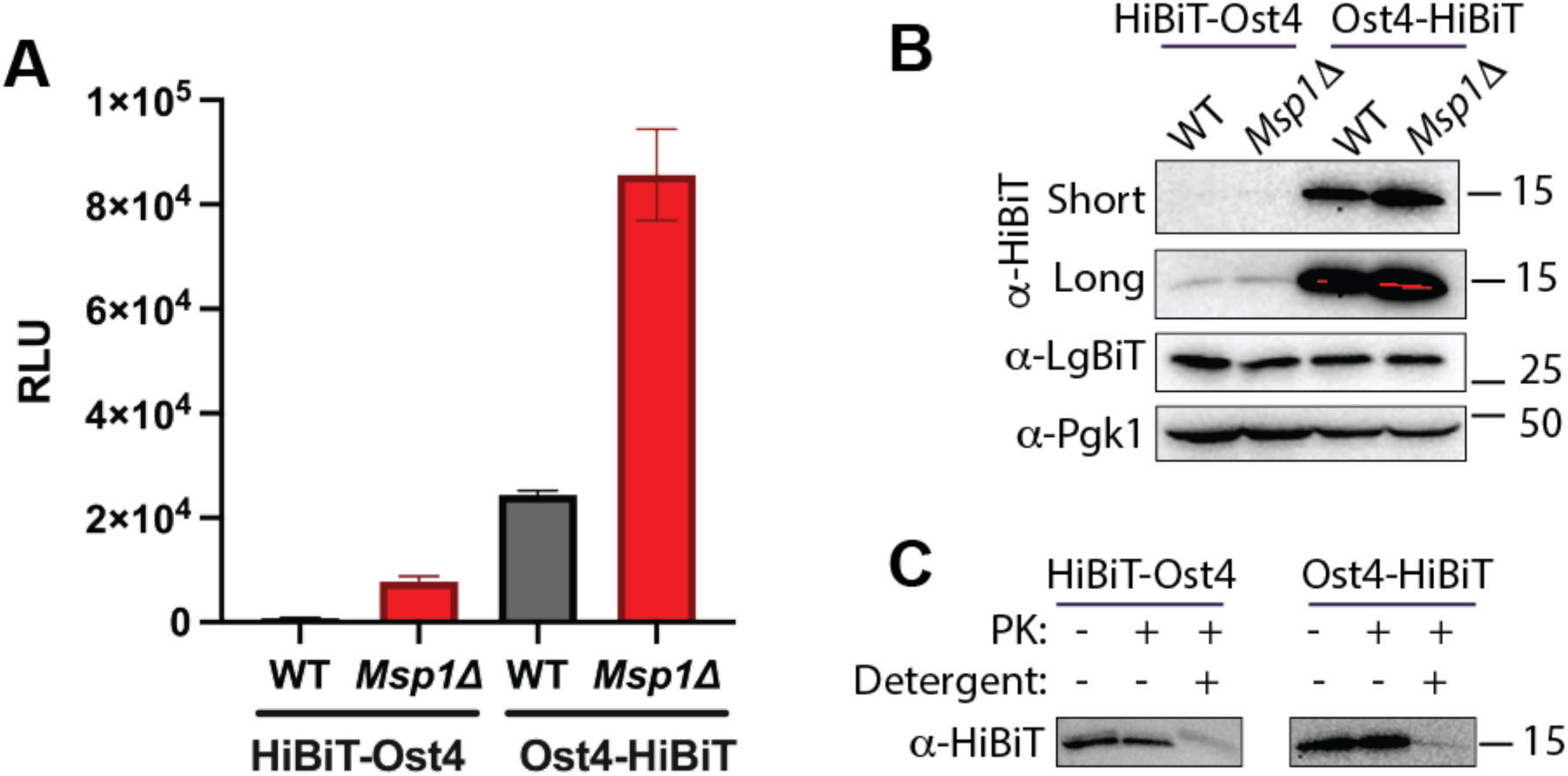
Ost4 adopts mixed topologies when mislocalized to mitochondria and both orientations are extracted by Msp1. A) Split luciferase assay shows an increase in luminescence in *Msp1*Δ cells regardless of whether the HiBiT tag is at the N- or C-terminus of Ost4. Error bars are standard error of the mean from 3 replicates. B) Western blot showing equal expression of LgBiT and Ost4-HiBiT or HiBit-Ost4 in WT and *Msp1Δ* cells. C) Protease protection assay on isolated mitochondria shows that the HiBiT tag is protected from proteolysis unless the membrane is solubilized by detergent.

To validate these surprising results, we performed a protease protection assay. *Msp1Δ* cells were transformed with a plasmid expressing Ost4-HiBiT or HiBiT-Ost4. Mitochondria were isolated and exposed proteins were digested with proteinase K. Consistent with the results of the luminescence assay, the C-terminal HiBiT is protected by the mitochondrial membrane (**Figure 4C**). We conclude that mislocalized Ost4 can adopt mixed topologies on the OMM, with either the N or C-terminus in the IMS.

### Msp1 can process substrates that fail to reconstitute into the membrane

Our cell-based assay shows an increase in the mitochondrial concentration of Ost4, regardless of whether the HiBiT tag is at the N or C-terminus, suggesting that Msp1 can extract Ost4 in both the N to C and C to N direction. To provide direct biochemical evidence that Msp1 extracts Ost4 regardless of topology in the OMM, we used our previously established *in vitro* system with fully purified components(Fresenius & Wohlever, 2021; Fresenius *et al*, 2023). This assay involves reconstituting purified substrate into proteoliposomes, performing a “preclearing” step to remove non-reconstituted substrate by adding the TMD binding chaperones SGTA and calmodulin, and then testing for Msp1-dependent extraction using a split luciferase assay (**Figure 5A**).

**Figure 5:**
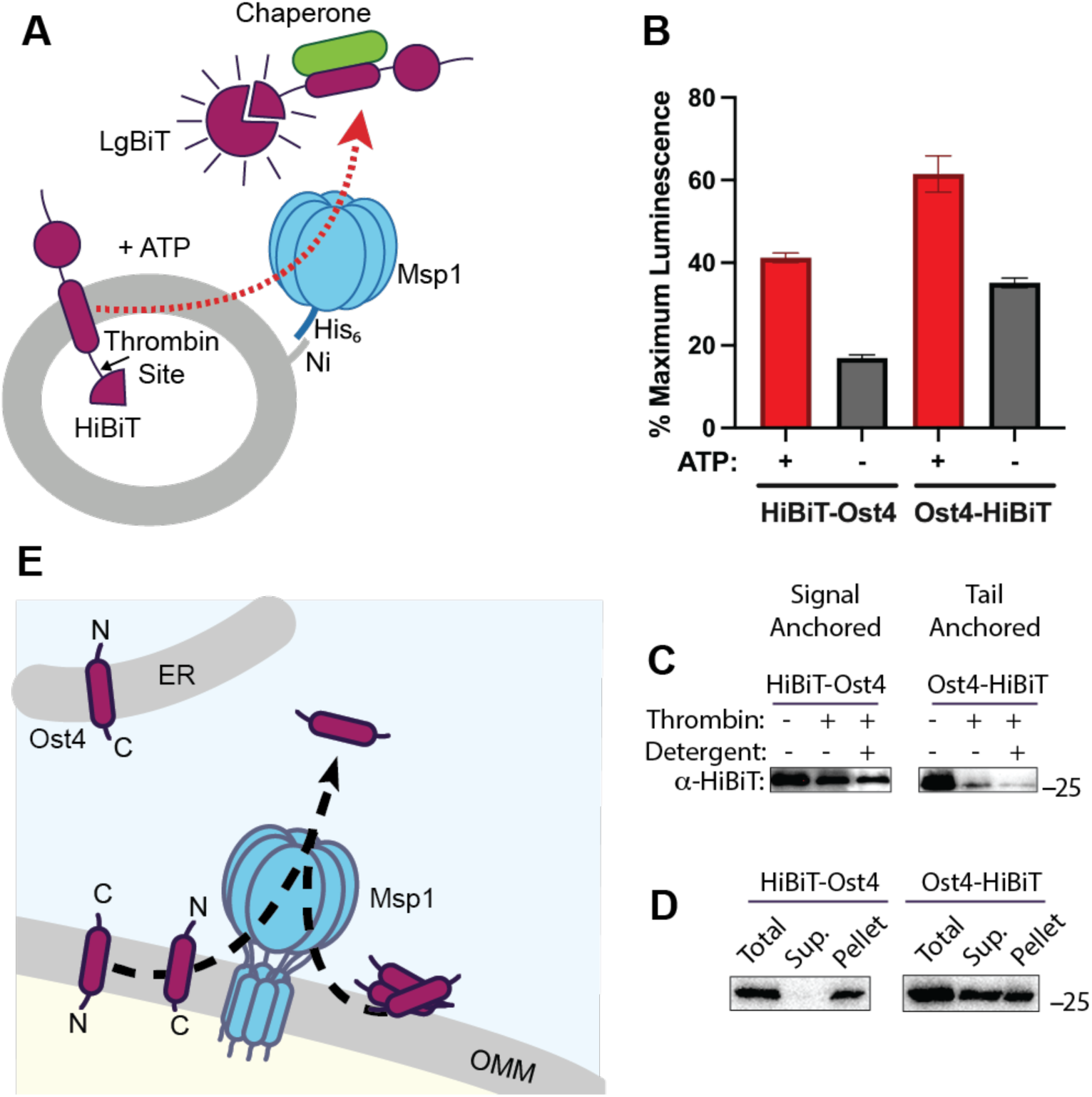
Msp1 processes Ost4 *in vitro*. A) Diagram of in vitro extraction assay. Substrate extraction is monitored by an increase in luminescence. B) In vitro extraction assay shows ATP dependent processing of both HiBiT-Ost4 and Ost4-HiBiT. C) Protease protection assay shows the majority of Ost4-HiBiT is not properly integrated in the liposome. Error bars are standard error of the mean from 3 replicates. D) Carbonate extraction assay on pre-cleared proteoliposomes shows that HiBiT-Ost4 is an integral membrane protein whereas the majority of Ost4-HiBiT is not fully integrated in the lipid bilayer. E) Model for Msp1 mediated extraction of Ost4. Ost4 normally resides in the ER with the N-terminus facing the ER lumen. Ost4 that is mistargeted to the OMM can have either terminus facing the IMS. Msp1 extracts Ost4 in both orientations. Msp1 can also process substrates that fail to integrate into the lipid bilayer.

We have previously demonstrated that the folded SUMO domain in our model substrate can bias substrate orientation during reconstitution into preformed liposomes (Fresenius *et al*, 2023; Wohlever *et al*, 2017; Fresenius & Wohlever, 2021). We therefore generated model substrates consisting of HiBiT-Ost4-SUMO and SUMO-Ost4-HiBiT to replicate signal anchored and tail-anchored orientations, respectively, of Ost4. The recombinantly expressed and purified substrates were then reconstituted into liposomes mimicking the lipid content of the OMM(Fresenius *et al*, 2023). We observed ATP dependent increase in luminescence with both the signal anchored and tail-anchored versions of Ost4 (**Figure 5B**).

We next attempted to validate substrate orientation with a protease protection assay. A thrombin protease site was engineered between the TMD and the HiBiT tag in the substrate. If the substrate is properly oriented with the thrombin cleavage site sequestered in the lumen of the liposome, then no substrate cleavage will be observed because thrombin cannot cross the lipid bilayer. Conversely, addition of detergent will solubilize the liposome, allowing thrombin to access the cleavage site, resulting in HiBiT peptide that is not resolvable by SDS PAGE/western blot. Our protease protection assay showed unexpected results, with signal anchored HiBiT-Ost4 having a population that is protease resistant, even in the presence of detergent, whereas the vast majority of tail-anchored Ost4-HiBiT is protease sensitive, even in the absence of detergent (**Figure 5C**). Numerous attempts to optimize reconstitution conditions or pre-clearing of unincorporated substrate failed to significantly improve results.

While it is formally possible that the majority of Ost4-HiBiT is inserted with the reverse topology, with the folded SUMO domain in the lumen and the small HiBiT tag facing bulk solvent, a more parsimonious explanation is the majority of the substrate failed to reconstitute into the liposome and is instead aggregated on the surface of the liposome where it is unable to interact with chaperones during the “preclearing” step. This would also explain the high levels of ATP-independent signal in the assay (**Figure 5B**). To test this hypothesis, we performed a carbonate extraction assay on the reconstituted, pre-cleared material. We observed that the signal anchored HiBiT-Ost4 construct was carbonate resistant, suggesting that it is fully integrated in the bilayer (**Figure 5D**). Conversely, the tail-anchored Ost4-HiBiT construct was mostly carbonate sensitive, suggesting a significant proportion of the population is not an integral membrane protein following reconstitution. Combined with the *in vivo* data (**Figure 4**), we conclude that, at least *in vitro*, Msp1 can process Ost4 regardless of whether it is a peripheral or integral membrane protein (**Figure 5E**).

## Discussion

AAA+ proteins are ubiquitous molecular motors, but identification of translocated substrates is challenging due to the transient nature of the interactions. While mutating the Walker B motif is effective at stabilizing substrate interactions with AAA+ proteins, inactivation of the enzyme can lead to undesired phenotypes. By fusing Msp1 to the inactivated protease domain of Yme1, we generated an Msp1-protease chimera that retains full Msp1 functionality but was also able to trap substrates better than WT Msp1. This protease chimera represents a critical advance in the study of AAA+ proteins, providing a powerful and versatile platform to dissect transient interactions without compromising catalytic activity. Its utility extends beyond Msp1, offering a broadly applicable strategy for unraveling the dynamic mechanisms of AAA+ motor proteins across diverse cellular pathways.

We performed mass spectrometry analysis with the Msp1-protease chimera under both standard conditions and mitochondrial import stress generated via CCCP treatment. After filtering the dataset to remove non-specific interactors, we obtained a list of 55 candidate substrates and validated that the proteins Ost4 and Phb1 specifically interact with Msp1. Interestingly, both Ost4 and Phb1 proteins have a single TMD at the N-terminus of the protein. All previously identified Msp1 substrates are tail-anchored proteins, with the C-terminus of the substrate in the IMS. Using an *in vivo* assay, we demonstrate that Msp1 can extract Ost4. This suggests that Msp1 can translocate substrates in both the N to C direction and the C to N direction, akin to other AAA+ proteins, such as ClpX(Barkow *et al*, 2009).

Although Ost4 is classified as a signal anchored protein, at only 36 residues long, it is likely inserted into the membrane post-translationally. While the initial characterization of Ost4 topology clearly shows that Ost4 is functional as a signal anchored protein, these experiments involved fusing an additional 128 kDa to the C-terminus of the 3.9 kDa Ost4 protein, which prevents tail-anchored topology(Kim *et al*, 2003). Our *in vivo* split luciferase assay overcomes this technical challenge as the 11 residue HiBiT sequence allows Ost4 to still be classified as a tail-anchored protein(Hall *et al*, 2012). Using this approach, we show that mislocalized Ost4 adopts mixed topologies in the OMM, with either the N or C-terminus in the IMS. It will be interesting to see if mixed topology is a general feature of proteins mislocalized to the OMM or is unique to small proteins such as Ost4, which lack a folded cytosolic domain to bias orientation in the lipid bilayer. How Ost4 adopts a signal anchored orientation in the ER in unclear and will be another interesting area of future investigation.

Attempts to visualize Msp1-dependent extraction of Ost4 *in vitro* were hampered by poor reconstitution of Ost4 into liposomes. Despite the low reconstitution efficiency, we observed ATP-dependent processing of Ost4. Combined with results of the carbonate extraction assay, this demonstrates that, at least *in vitro*, Msp1 can process peripheral membrane proteins. As previous work has shown that Msp1 recognizes exposed hydrophobic residues(Fresenius *et al*, 2023; Li *et al*, 2019), it is tempting to speculate that Msp1 can also function as a disaggregase for proteins on the OMM. This proposal is particularly appealing because protein aggregates are known to accumulate on the mitochondrial surface (Liu *et al*, 2023) and AAA+ proteins commonly serve as Disaggregases (Wohlever *et al*, 2013; Cupo & Shorter, 2020; Shorter & Southworth, 2019). Demonstrating Msp1 disaggregation activity *in vivo* will depend on identification of physiological substrates and our data set provides a robust starting point for this task.

In summary, we have developed a generalizable strategy for stabilizing the transient interaction of substrates with AAA+ proteins that does not require inactivation of ATPase activity, expanded the Msp1 substrate repertoire to include multiple substrates with an N-terminal TMD, demonstrated that mitochondrial localized Ost4 adopts mixed topologies, and shown that Msp1 can extract substrates in both the N to C and C to N direction. Together, this work expands our understanding of protein mistargeting to the OMM and the role of Msp1 in proteostasis.

## Methods

**Table 1:**
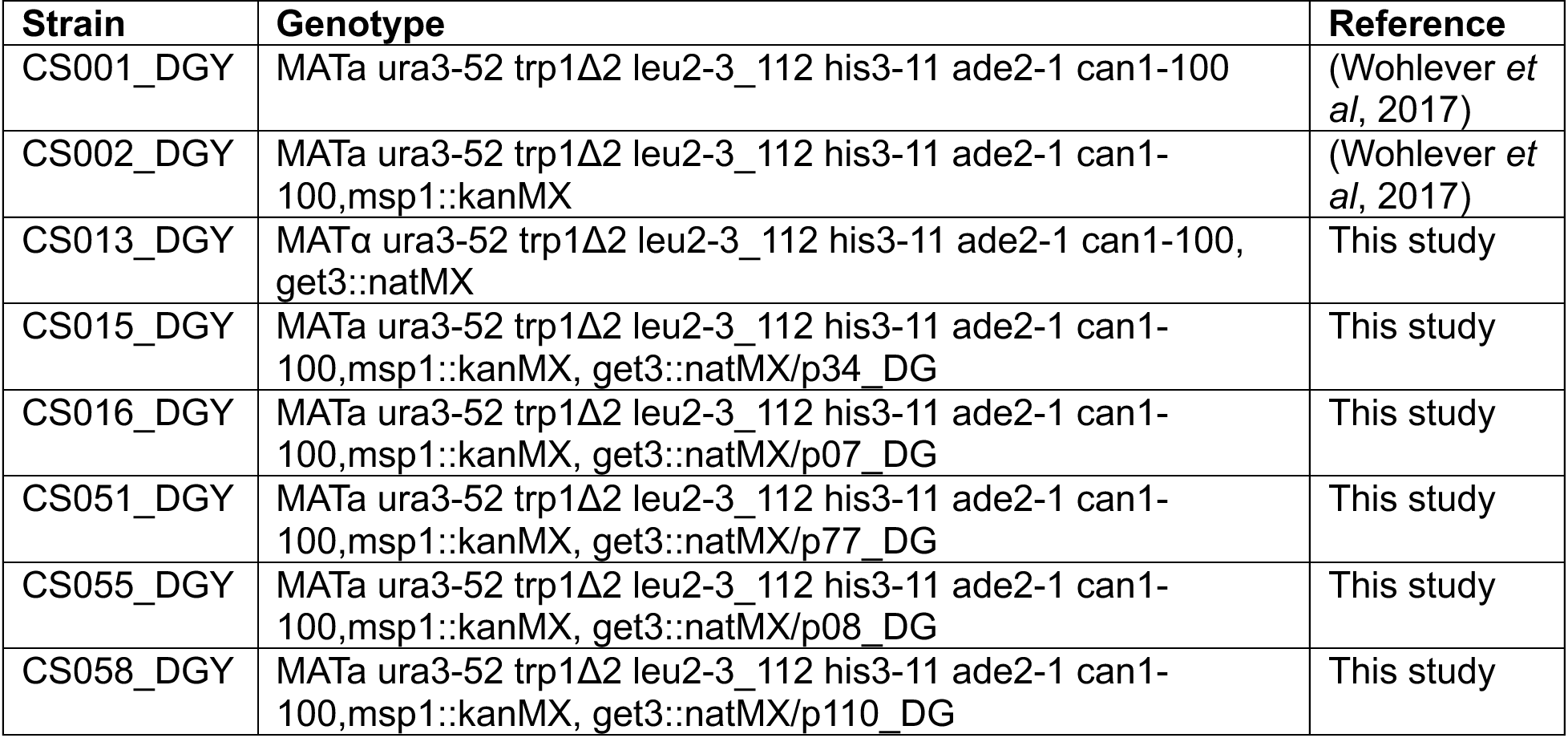
Strains used in this study.

**Table 2:**
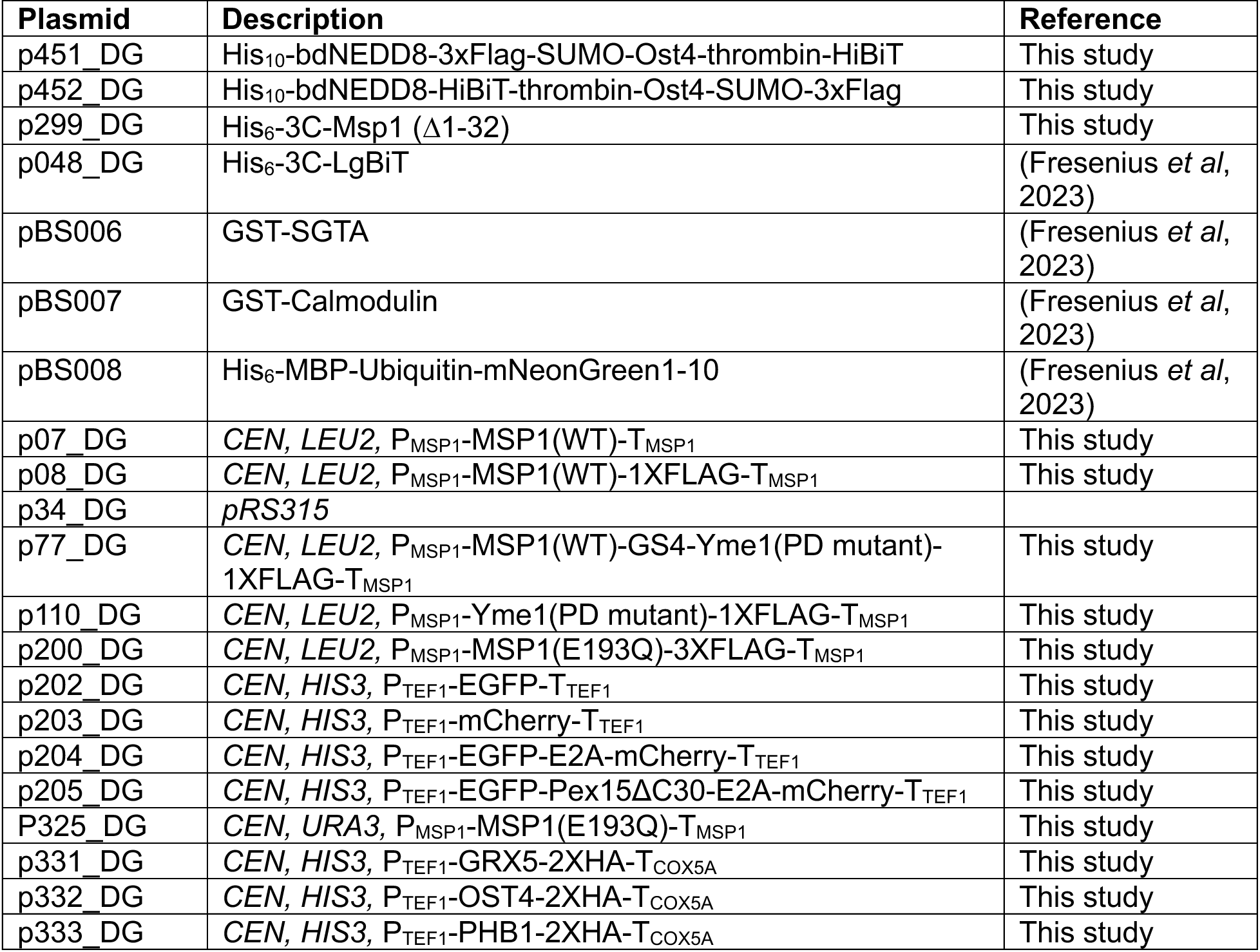
Plasmid used in this study.

| Plasmid | Description | Reference |
| --- | --- | --- |
| p451_DG | His<sub>10</sub>-bdNEDD8-3xFlag-SUMO-Ost4-thrombin-HiBiT | This study |
| p452_DG | His<sub>10</sub>-bdNEDD8-HiBiT-thrombin-Ost4-SUMO-3xFlag | This study |
| p299_DG | His<sub>6</sub>-3C-Msp1 (Δ1-32) | This study |
| p048_DG | His<sub>6</sub>-3C-LgBiT | (Fresenius *et al*, 2023) |
| pBS006 | GST-SGTA | (Fresenius *et al*, 2023) |
| pBS007 | GST-Calmodulin | (Fresenius *et al*, 2023) |
| pBS008 | His<sub>6</sub>-MBP-Ubiquitin-mNeonGreen1-10 | (Fresenius *et al*, 2023) |
| p07_DG | CEN, LEU2, P<sub>MSP1</sub>-MSP1(WT)-T<sub>MSP1</sub> | This study |
| p08_DG | CEN, LEU2, P<sub>MSP1</sub>-MSP1(WT)-1XFLAG-T<sub>MSP1</sub> | This study |
| p34_DG | pRS315 |  |
| p77_DG | CEN, LEU2, P<sub>MSP1</sub>-MSP1(WT)-GS4-Yme1(PD mutant)-1XFLAG-T<sub>MSP1</sub> | This study |
| p110_DG | CEN, LEU2, P<sub>MSP1</sub>-Yme1(PD mutant)-1XFLAG-T<sub>MSP1</sub> | This study |
| p200_DG | CEN, LEU2, P<sub>MSP1</sub>-MSP1(E193Q)-3XFLAG-T<sub>MSP1</sub> | This study |
| p202_DG | CEN, HIS3, P<sub>TEF1</sub>-EGFP-T<sub>TEF1</sub> | This study |
| p203_DG | CEN, HIS3, P<sub>TEF1</sub>-mCherry-T<sub>TEF1</sub> | This study |
| p204_DG | CEN, HIS3, P<sub>TEF1</sub>-EGFP-E2A-mCherry-T<sub>TEF1</sub> | This study |
| p205_DG | CEN, HIS3, P<sub>TEF1</sub>-EGFP-Pex15ΔC30-E2A-mCherry-T<sub>TEF1</sub> | This study |
| P325_DG | CEN, URA3, P<sub>MSP1</sub>-MSP1(E193Q)-T<sub>MSP1</sub> | This study |
| p331_DG | CEN, HIS3, P<sub>TEF1</sub>-GRX5-2XHA-T<sub>COX5A</sub> | This study |
| p332_DG | CEN, HIS3, P<sub>TEF1</sub>-OST4-2XHA-T<sub>COX5A</sub> | This study |
| p333_DG | CEN, HIS3, P<sub>TEF1</sub>-PHB1-2XHA-T<sub>COX5A</sub> | This study |
| p337_DG | *CEN, HIS3, P<sub>TEF1</sub>-RIP1-2XHA-T<sub>COX5A</sub>* | This study |
| p338_DG | *CEN, HIS3, P<sub>TEF1</sub>-VTC3-2XHA-T<sub>COX5A</sub>* | This study |

### Strain construction

Strains and plasmids used in the study are described in **Tables 1** and **2**. For complementation assays using different Msp1 constructs in *S. cerevisiae*, *Msp1Δ, Get3Δ* W303-1 yeast were generated by integrating Msp1::KanMX and Get3::NatMX using homologous recombination individually in WT 303-1 strain. The diploid strain (Msp1::KanMX and Get3::NatMX) was constructed by mating CS002_DGY *MATα* (Msp1::KanMX) with CS0013_DGY *MATa* (Get3::NatMX). CS002_DGY *MATα* (Msp1::KanMX) was also transformed with different Msp1 constructs cloned in *CEN* and *LEU2* vector with the flanking 281 bp upstream and 261 bp downstream as the respective promoter and terminator. Diploids were sporulated on a 2% potassium acetate plate and tetrad dissection was carried out on YPAD plates. Resulting tetrads were selected on plates containing G418 (200 μg/ml), NAT (200 μg/ml) and lacking leucine amino acid. Tetrads were further replica plated onto leucine deficient SD plate containing G418 and NAT.

### Construct design

Ost4-SUMO constructs and soluble Msp1 for *in vitro* extraction assay were generated by gene synthesis and cloned into a Golden Gate domesticated pET28b vector(Gaur & Wohlever, 2025). The protease chimera was generated by Gibson Assembly with Msp1 and Yme1 sequences isolated from genomic DNA. Potential Msp1 substrates were amplified from genomic DNA and cloned into centromeric plasmids via golden gate cloning(Agmon *et al*, 2015).

### Yeast based assays

#### Glycerol complementation assay

Briefly, haploid W303-1 cells containing appropriate Msp1 plasmid cloned in *CEN* and *LEU2* vector with the flanking 281 bp upstream and 261 bp downstream as the respective promoter and terminator were grown overnight in SD-LEU medium with 2% glucose. Cultures were diluted to 0.02 OD_600nm_ in fresh SD-LEU media until the mid-log phase. Cultures were washed 3x with sterile water and diluted to an OD_600nm_ of 1. Samples were serially diluted 5x and then spotted onto SD-LEU plates containing either 2% glucose or 2% glycerol and grown at 30 °C. Images are representative of N > 2 trials.

#### Pex15 overexpression

Haploid W303-1 cells containing the appropriate Msp1 plasmid cloned in *CEN* and *LEU2* vector were transformed with Pex15 cloned under GAL1 promoter in a CEN and URA vector. The transformants were grown overnight in SD-LEU-URA medium with 2% glucose. Cultures were further diluted at 0.02 OD_600nm_ in fresh SD-LEU-URA media until the OD_600nm_ between 1.5-2. Cultures were washed 3x with sterile water and diluted to 1 OD_600nm_. Samples were serially diluted 5x and then spotted onto SD and SGAL plates lacking leucine and uracil amino and grown at 30 °C. SD and SGAL plates were imaged after 2 and 5 days respectively. Images are representative of N > 2 trials.

#### Split luciferase assay

WT W303-1 and *Msp1Δ* W303-1yeast strains were co-transformed with substrates (HiBiT-Ost4-, Ost4-HiBiT, HiBiT-Phb1, and Phb1-HiBiT) cloned in CEN and HIS vector under strong TEF promoter and mito-LgBiT cloned in CEN and URA vector under RNR1 promoter. Transformants were grown overnight in SD-His-Ura media, followed by diluting the grown cultures at 0.02 OD_600nm_ in fresh SD-His-Ura media until OD_600nm_ = 2.0 and 40 µL of cells were transferred to a 96 well plate. The reaction was incubated at 30°C with Nano-Glo Live Cell assay reagent (N2012, Promega) and the interaction was monitored as an increase in luminescence with time in a Spectramax iD5 plate reader (Molecular Devices). For substrates under control of the GAL1 promoter (Pex15-HiBiT, Bos1-HiBiT, Bos1-6AG-HiBiT), constructs were cloned in CEN and URA vector and yeast cells were subsequently induced in SGAL-HIS-URA media for 8 hours.

Samples were washed 2X with sterile water followed by dilution to OD_600nm_ = 2.0 in sterile water and then luminescence was monitored as described above.

### Immunoprecipitations

For immunoprecipitation, cells were resuspended in IP Buffer (50 mM Hepes⋅KOH pH 7.5, 50 mM Potassium acetate, 2 mM Magnesium acetate, 200 mM sorbitol, 0.1% digitonin, 1x complete protease inhibitors-EDTA-free, and 1 mM PMSF), and lysed by mechanical disruption using glass beads. The resulting lysate was solubilized with 0.5% digitonin for 1 hour at 4°C. Lysate was clarified by centrifuging at 12000g for 15 minutes. The resulting supernatant (2.5 mg protein) was incubated for 1 hour at 4°C with anti-FLAG antibody resin (A220, Sigma). The beads were washed 4 times with IP buffer containing 0, 150, 300, and 500 mM NaCl. The bound protein was eluted using 1mg/ml FLAG peptide. Immunoprecipitated proteins were further separated on SDS-PAGE and immunoblotted with different antibodies.

### Western blots

Cells grown in liquid media were harvested by centrifugation. To facilitate the permeabilization of the cellular wall, the cells were treated with 0.2N NaOH for a duration of 10 minutes while maintained on ice. Subsequently, the treated cells were subjected to centrifugation at 13000 rpm for 1 minute, followed by the solubilization and denaturation of proteins in 1X SDS loading dye at 95°C for 5 minutes. The proteins were separated by SDS-PAGE, transferred onto PVDF membranes, and probed using the appropriate antibodies. The primary antibodies used in the study were as follows: anti-FLAG (F7425, Sigma), Anti-GFP (MA5-15256, Thermo Fisher Scientific), Anti-HiBiT (N720A, Promega), Anti-LgBiT (N710A, Promega), Anti-HA (26183, Invitrogen), Anti-Pgk1 (459250, Thermo Scientific), Anti-mouse secondary (31430, Invitrogen), and Anti-Rabbit secondary (SA00001-2, Proteintech). Primary antibodies were used at dilution of 1:5000 and incubated with an immunoblot for 1 hour followed by 40 minutes incubation with secondary antibodies at room temperature.

### Protease protection assay on isolated mitochondria

Total of 50 µg of crudely purified mitochondria were incubated under various conditions to assess the orientation of different substrates. Briefly mitochondria were incubated for 10 minutes in iso-osmolar SEM buffer (10 mM MOPS/KOH (pH 7.2), 250 mM sucrose, 1 mM EDTA) in the presence and absence of 1% Triton X-100. This is followed by addition of 20 μg/ml of Proteinase-K. Reaction is halted by addition of 4mM PMSF. Mitochondria is pelleted by centrifuging at 12000g for 15 minutes. Resulting pelleted fraction is denatured in 1X SDS loading dye at 95°C for 5 minutes. Denatured samples are separated on 15% SDS-PAGE followed by immunoblotting with appropriate antibodies.

### Mass spectrometry and data analysis

Mass spectrometry (MS) was performed at the University of Michigan chemistry mass spectrometry facility. Elution from anti-Flag IP were prepared for mass spectrometry via disulfide reduction, aklylation, and trypsin digestion at 40° C overnight. The samples purified with a zirconium dioxide (ZrO_2_) column and reconstituted in 2% Acetonitrile with 0.1% formic acid. Digested peptides were separated on a C18 column (Acclaim PepMap 100 C18 HPLC column, #164942) and analyzed on the nanoUPLC (Ultimate 3500) linked to Orbitrap Fusion Lumos MS via ESI(+) online. Protein identification was performed by Higher Energy Collisional Dissociation MS-MS in Data Dependent Acquisition mode and then analyzed with Protein Discoverer (v 2.2) against UniProt-YEAST database. Common contaminants, M-oxidation, peptide n-terminal carbamylation, protein N-terminal acetylation, and fixed C-cabamidomethylation were included in the search. A maximum of five missed trypsin cleavages were allowed. A false discovery rate filter of 1% was applied for peptide and protein IDs.

### Soluble Protein Purification Δ1-32 Msp1

Soluble Msp1 was purified as previously described(Fresenius *et al*, 2023). Plasmids were transformed into *E. coli* LOBSTR BL21(DE3) containing a pRIL plasmid and expressed in terrific broth at 37°C until an OD_600nm_ of 0.6–0.8, cultures were induced with 0.25 mM IPTG and grown at 16° C for 16 hours(Andersen *et al*, 2013). Cells were harvested by centrifugation, and resuspended in Msp1 Lysis Buffer (20 mM Tris pH 7.5, 200 mM potassium acetate, 20 mM imidazole, 0.01 mM EDTA, 1 mM DTT) supplemented with 0.05 mg/mL lysozyme (Sigma), 1 mM phenylmethanesulfonyl fluoride (PMSF) and 500 U of universal nuclease (Pierce). Sample was lysed by sonication and insoluble material was pelleted by centrifugation for 30 min at 4°C at 18,500 × g. The supernatant was purified by Ni-NTA affinity chromatography (Qiagen) on a gravity column. Ni-NTA resin was washed with 10 column volumes (CV) of Msp1 Lysis Buffer and then 10 CV of Wash Buffer (20 mM Tris pH 7.5, 200 mM potassium acetate, 30 mM imidazole, 0.01 mM EDTA, 1 mM DTT) before elution with Elution Buffer (20 mM Tris pH 7.5, 200 mM potassium acetate, 250 mM imidazole, 0.01 mM EDTA, 1 mM DTT).

The protein was further purified by size exclusion chromatography (SEC) (Superdex 200 Increase 10/300 GL, Cytiva) in 20 mM Tris pH 7.5, 200 mM KAc, 1 mM DTT. Peak fractions were pooled, concentrated to 5–10 mg/mL in a 30 kDa MWCO Amicon Ultra centrifugal filter (Pierce) and aliquots were flash-frozen in liquid nitrogen and stored at –80°C. Protein concentrations were determined by A280 using a calculated extinction coefficient (Expasy).

#### SGTA and Calmodulin

GST-SGTA and GST-calmodulin were expressed, pelleted, and lysed as described above for soluble Msp1 constructs, except that SGTA Lysis Buffer was used (50 mM HEPES pH 7.5, 150 mM NaCl, 0.01 mM EDTA, 1 mM DTT, 10% glycerol). Following removal of insoluble material by centrifugation, the supernatant was purified by glutathione affinity chromatography (Thermo Fisher) on a gravity column. Resin was washed with 20 CV of SGTA Lysis Buffer and then eluted with 3 CV of SGTA Lysis Buffer supplemented with 10 mM reduced glutathione. The protein was further purified by SEC (Superdex 200 Increase 10/300 GL, Cytiva) in SGTA FPLC Buffer (20 mM Hepes pH 7.5, 100 mM NaCl, 0.1 mM TCEP). Peak fractions were pooled, concentrated to 20 mg/mL in a 30 kDa (Calmodulin) or 50 kDa (SGTA) MWCO Spin Concentrator (Amicon) and aliquots were flash-frozen in liquid nitrogen and stored at –80°C. Protein concentrations were determined by A280 using a calculated extinction coefficient (Expasy).

#### LgBiT and MBP-Ubiquitin

LgBiT and MBP-Ubiquitin were expressed as described above for soluble Msp1 constructs except LgBiT Lysis Buffer was used (20 mM Tris pH 7.5, 200 NaCl, 1 DTT, 0.01 EDTA, 20 mM imidazole). Following centrifugation, the supernatant was purified by Ni-NTA affinity chromatography (Qiagen) on a gravity column. Ni-NTA resin was washed with 10 column volumes (CV) of LgBiT Lysis Buffer and then 10 CV of LgBiT Wash Buffer (20 mM Tris pH 7.5, 200 NaCl, 1 DTT, 0.01 EDTA, 50 mM imidazole) before elution with LgBiT Lysis Buffer (20 mM Tris pH 7.5, 200 NaCl, 1 DTT, 0.01 EDTA, 250 mM imidazole). The protein was further purified by size exclusion chromatography (SEC) (Superdex 200 Increase 10/300 GL, Cytiva) in LgBiT FPLC Buffer (20 mM Tris pH 7.5, 200 mM NaCl, 1 mM DTT). Peak fractions were pooled, concentrated to 5–15 mg/mL in a 30 kDa MWCO Amicon Ultra centrifugal filter (Pierce) and aliquots were flash-frozen in liquid nitrogen and stored at –80°C. Protein concentrations were determined by A280 using a calculated extinction coefficient (Expasy).

### Membrane Protein Purification Ost4 -SUMO constructs

The SUMO-Ost4 fusion constructs were expressed, pelleted, and lysed as described above for soluble Msp1 constructs, except that SUMO TMD Lysis Buffer was used (50 mM Tris pH 7.5, 300 mM NaCl, 10 mM MgCl_2_, 25 mM Imidazole, 10% glycerol). Lysate was solubilized by the addition of n-dodecyl-β-D-maltoside (DDM) to a final concentration of 1% and rocked at 4°C for 1 h. Lysate was cleared by centrifugation for at 4°C for 1 h at 35,000 x g and purified by Ni-NTA affinity chromatography.

Ni-NTA resin was washed with 5 column volumes of SUMO TMD Wash Buffer 1 (50 mM Tris pH 7.5, 500 mM NaCl, 10 mM MgCl_2_, 25 mM imidazole, 5 mM β-mercaptoethanol (BME), 10% glycerol, 0.1% DDM). Resin was then washed with 5 CV of SUMO TMD Wash Buffer 2 (50 mM Tris pH 7.5, 300 mM NaCl, 10 mM MgCl_2_, 40 mM imidazole, 5 mM BME, 10% glycerol, 0.1% DDM), 10 CV of SUMO TMD Wash Buffer 3 (50 mM Tris pH 7.5, 150 mM NaCl, 10 mM MgCl_2_, 50 mM imidazole, 5 mM BME, 10% glycerol, 0.1% DDM) and then eluted with 3 CV of SUMO TMD Elution Buffer (50 mM Tris pH 7.5, 150 mM NaCl, 10 mM MgCl_2_, 250 mM imidazole, 5 mM BME, 10% glycerol, 0.1% DDM)

The protein was further purified by size exclusion chromatography (SEC) (Superdex 200 Increase 10/300 GL, Cytiva) in SUMO TMD FPLC Buffer (50 mM Tris pH 7.5, 150 mM NaCl, 10 mM MgCl_2_, 5 mM BME, 10% glycerol, 0.1% DDM). Peak fractions were pooled, aliquots were flash-frozen in liquid nitrogen and stored at –80°C. Protein concentrations were determined by A_280_ using a calculated extinction coefficient (Expasy).

### Liposome preparation

Liposomes mimicking the lipid composition of the yeast OMM were prepared as described previously(Fresenius *et al*, 2023; Fresenius & Wohlever, 2021). A lipid film was prepared by mixing chloroform stocks of the following lipids: chicken egg phosphatidyl choline (Avanti 840051C), chicken egg phosphatidyl ethanolamine (Avanti 840021C), bovine liver phosphatidyl inositol (Avanti 840042C), synthetic DOPS (Avanti 840035C), synthetic TOCL (Avanti 710335C), and 1,2-dioleoyl-*sn*-glycero-3-[*N*-((5-amino-1-carboxypentyl)iminodiacetic acid)succinyl] Nickel salt (Avanti 790404) at a 48:28:10:8:4:2 molar ratio with 1 mg of DTT. Chloroform was evaporated under a gentle steam of nitrogen and then left on a vacuum (<1 mTorr) overnight. Lipid film was fully resuspended in Liposome Buffer (50 mM HEPES KOH pH 7.5, 15% glycerol, 1 mM DTT) to a final concentration of 20 mg/mL and then subjected to five freeze-thaw cycles with liquid nitrogen. Liposomes were extruded 15 times through a 100 nm filter at 60°C, distributed into single-use aliquots, and flash-frozen in liquid nitrogen.

### Reconstitution into Liposomes

For extraction assays proteoliposomes were prepared by mixing 2.5 μM of the appropriate Ost4-SUMO construct, and 2 mg/mL of Nickel liposomes in Reconstitution Buffer(Fresenius *et al*, 2023; Fresenius & Wohlever, 2021). Detergent was removed by adding 25 mg of biobeads and rotating the samples for 16 hours at 4°C. Material was removed from biobeads. Unincorporated TA protein was pre-cleared by incubating the reconstituted material with excess (5 μM) GST-SGTA and GST-calmodulin and passing over a glutathione spin column (Pierce #16103); the flow through was collected and used immediately for dislocation assays.

### Extraction Assay

Extraction assays were performed as described previously(Fresenius *et al*, 2023; Fresenius & Wohlever, 2021). Each reaction contained 30 μL of pre-cleared proteoliposomes, 5 μM GST-SGTA, 5 μM GST-calmodulin, 3 μM Msp1, 8 μM MBP-Ubiquitin, 1 mg/mL bovine serum albumin (sigma), and 80 mM ATP and the final volume was adjusted to 105 μL with Extraction Buffer (50 mM HEPES KOH pH 7.5, 200 mM potassium acetate, 7 mM magnesium acetate, 2 mM DTT, 0.1 μM calcium chloride). Samples were incubated at 30°C for 30 minutes and then transferred to ice. Samples were transferred to polycarbonate centrifuge tubes (Beckman-Coulter #343776) and centrifuged at 100,000 x *g* in S100-AT3 Fixed-Angle Rotor (Thermo Fisher) for 30’ at 4°C. The top 20 ul was collected and combined with 1.5 μM LgBiT and the final volume was adjusted to 50 μL with Extraction Buffer. After incubating at RT for 25 minutes, samples were transferred into white 96-well plates (Corning #07201204) and 20 μL of lytic furimazine (Promega #N3030) was added. Immediately plates were read at 480 nm wavelength to measure luminescence.

Percent substrate extracted was calculated by comparing to a standard curve containing 5 different concentrations of the same pre-cleared liposomes used in the extraction combined with Extraction Buffer (50 mM HEPES KOH pH 7.5, 200 mM potassium acetate, 7 mM magnesium acetate, 2 mM DTT, 0.1 μM calcium chloride). A total of 20 μL from each standard was then combined with 1.5 μM LgBiT and the final volume was adjusted to 50 μL with Extraction Buffer. After incubating at RT for 25 minutes, samples were plated onto white 96-well plates (Corning #07201204) and 20 μL of lytic furimazine (Promega #N3030) was added. Immediately plates were read at 480 nm wavelength to measure luminescence.

### Liposome Protease Protection Assay

Protease protection assay was carried out with 7 μL of pre-cleared liposomes, 2 U of thrombin protease, and 1% of Triton X-100 (where indicated) in a total volume of 10 μL(Fresenius *et al*, 2023). Samples were incubated at RT for 1 hour and then 2 μL of 100 mM PMSF was added. The samples were reverse quenched into 90 μL of boiling 1% SDS and incubated at 95°C for 10 minutes, followed by anti-HiBiT western blot.

### Liposome Carbonate Extraction Assay

Carbonate extraction assay was carried out as previously described(Wohlever *et al*, 2017). Briefly, substrate was reconstituted into liposomes and pre-cleared as described above. The sample was then diluted 2x with 100 mM sodium carbonate pH 11.5 and incubated on ice for 15 minutes. The membrane associated fraction was pelleted by spinning for 45 minutes at 100,000 x g in a S100-AT3 rotor. The pellet was resuspended in an equal volume of 1x SDS Loading Buffer.

## Data availability

All data is available upon request.

## Acknowledgements

The authors wish to thank members of the Wohlever for helpful discussions and feedback on the project. Mass spectrometry work was completed at the University of Michigan core facility.

## Funding

This work was supported by NIH grant R35GM137904 (MLW). The mass spec core at the University of Michigan. The Orbitrap Fusion Lumos is supported by the Office of the Director, National Institutes of Health under Award Number S100D021619.

## Author Contributions

Conceptualization: DG, MLW; Data curation: DG, BA, MLW; Formal analysis: DG, BA, MLW; Funding acquisition: MLW; Investigation: DG, BA; Methodology: DG, BA; Project administration: MLW; Resources: DG, BA, MLW; Software: Not applicable; Supervision: MLW; Validation: DG, BA; Visualization: DG, BA, MLW; Writing – original draft: DG, MLW; Writing – review & editing: DG, BA, MLW;

## Competing interests

The authors declare no competing interests

## Disclosures and competing interests

The authors declare no competing interests

## Supplemental Figures

**Figure S1:**
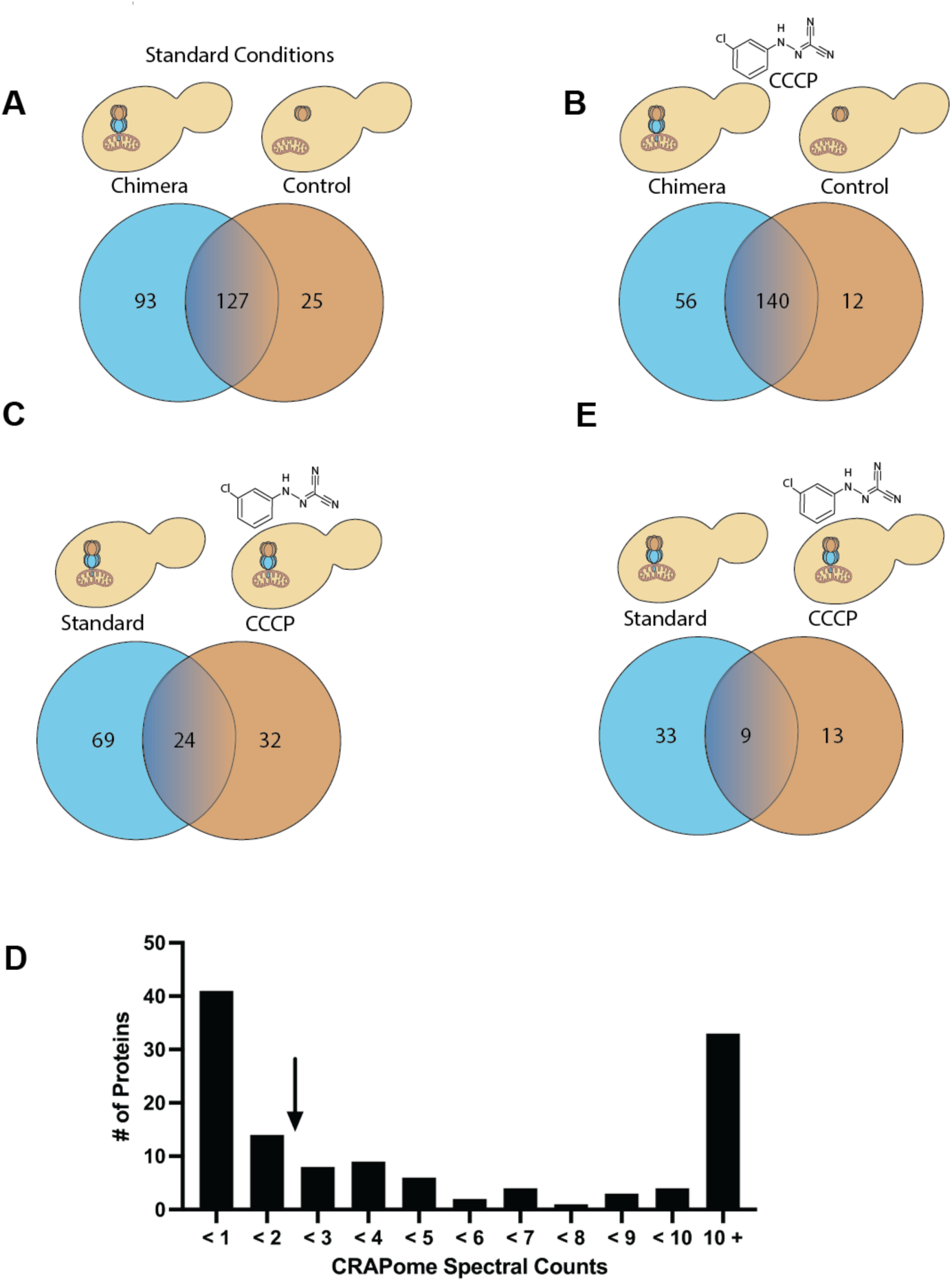
Mass spectrometry analysis of Msp1-Protease Chimera interactions. A) Venn Diagram showing number of interactors with Msp1-Protease Chimera and protease domain alone negative control for “standard conditions” of SD – Leu. B) As in A, but for cells treated with 20 μM CCCP for 2 h prior to immunoprecipitation. C) Venn diagram showing number of hits unique to the Msp1-Protease Chimera under standard conditions and after CCCP treatment. D) Histogram of candidate proteins from the combined standard and CCCP datasets binned by the number of spectral counts that also appear in the CRAPome database. The final list of candidate proteins are to the left of the arrow. E) Venn diagram as in C, but this time filtered to only include proteins with CRAPome spectral counts below 2.

**Table S1: Chimera mass spec results for standard conditions dataset**

**Table S2: Protease domain control mass spec results for standard dataset**

**Table S3: Chimera mass spec results for CCCP dataset**

**Table S4: Protease domain control mass spec results for CCCP dataset**

**Table S5: Combined, chimera specific hits filtered by CRAPome spectral counts**

## Notes

### Competing Interest Statement

The authors have declared no competing interest.

